# Preferential differential gene expression within the WC1.1^+^ γδ T cell compartment in cattle naturally infected with *Mycobacterium bovis*

**DOI:** 10.1101/2023.07.21.550071

**Authors:** Sajad A. Bhat, Mahmoud Elnaggar, Thomas J. Hall, Gillian P. McHugo, Cian Reid, David E. MacHugh, Kieran G. Meade

**Affiliations:** UCD School of Agriculture and Food Science, University College Dublin, Belfield, D04 V1W8, Dublin, Ireland; UCD Conway Institute of Biomolecular and Biomedical Research, University College Dublin, Belfield, D04 V1W8, Dublin, Ireland; Animal and Bioscience Research Department, Animal and Grassland Research and Innovation Centre, Teagasc, Grange, C15 PW93, Meath, Ireland; Department of Veterinary Medicine, College of Applied and Health Sciences, A’Sharqiyah University, Ibra, Oman; Department of Microbiology, Faculty of Veterinary Medicine, Alexandria University, Egypt; UCD Institute of Food and Health, University College Dublin, Belfield, D04 V1W8, Dublin, Ireland

**Keywords:** WC1^+^ γδ T cells, natural *Mycobacterium bovis*, bovine, NK receptors

## Abstract

Bovine tuberculosis (bTB), caused by infection with *Mycobacterium bovis*, continues to cause significant issues for the global agriculture industry as well as for human health. An incomplete understanding of the host immune response contributes to the challenges of control and eradication of this zoonotic disease. In this study, high-throughput bulk RNA sequencing (RNA-seq) was used to characterize differential gene expression in γδ T cells – a subgroup of T cells that bridge innate and adaptive immunity and have known anti-mycobacterial response mechanisms. γδ T cell subsets are classified based on expression of a pathogen-recognition receptor known as *Workshop Cluster 1* (WC1) and we hypothesised that bTB disease may alter the phenotype and function of specific γδ T cell subsets. Peripheral blood was collected from naturally *M. bovis*-infected (positive for single intradermal comparative tuberculin test (SICTT) and IFN-γ ELISA) and age- and sex-matched, non-infected control Holstein-Friesian cattle. γδ T subsets were isolated using fluorescence activated cell sorting (*n* = 10–12 per group) and high-quality RNA extracted from each purified lymphocyte subset (WC1.1^+^, WC1.2^+^, WC1^-^ and γδ^-^) was used to generate transcriptomes using bulk RNA-seq (*n* = 6 per group, representing a total of 48 RNA-seq libraries). Relatively low numbers of differentially expressed genes (DEGs) were observed between most cell subsets; however, 189 genes were significantly differentially expressed in the *M. bovis*-infected compared to the control groups for the WC1.1^+^ γδ T cell compartment (absolute log_2_ FC ≥ 1.5 and FDR *P*_adj._ ≤ 0.1). The majority of these DEGs (168) were significantly increased in expression in cells from the bTB+ cattle and included genes encoding transcription factors (*TBX21* and *EOMES*), chemokine receptors (*CCR5* and *CCR7*), granzymes (*GZMA, GZMM*, and *GZMH*) and multiple killer cell immunoglobulin-like receptor (KIR) proteins indicating cytotoxic functions. Biological pathway overrepresentation analysis revealed enrichment of genes with multiple immune functions including cell activation, proliferation, chemotaxis, and cytotoxicity of lymphocytes. In conclusion, WC1.1^+^ γδ T cells have been proposed as major regulatory cell subset in cattle, and we provide evidence for preferential differential activation of this specific subset in cattle naturally infected with *M. bovis*.

## INTRODUCTION

Bovine tuberculosis (bTB), caused by *Mycobacterium bovis* is endemic in many countries and has significant economic and animal welfare impacts across global agricultural systems (1). Additionally, *M. bovis* is a member of the *Mycobacterium tuberculosis* complex (MTBC), which can infect multiple species including humans and livestock making it a zoonotic pathogen and a significant threat to public health (2). The reasons for a failure to eradicate bTB are multifactorial (3) but scientific understanding of *M. bovis* persistence has been limited in part due to the complexity of host-pathogen interactions that occur under conditions of natural infection. The immune system in humans as well as in livestock is key to disease resistance (4) and suboptimal or dysregulated immunity is thought to contribute to bTB disease susceptibility, progression to clinical disease, and the failure of current generation diagnostics to detect all truly infected cattle. Addressing this important knowledge gap is critical toward achieving the ultimate goal of bTB eradication.

As an intracellular pathogen, immunological research on *M. bovis* has logically concentrated on the cell-mediated arm of the immune response, and in particular on the CD4^+^ helper T cell subset (5). As major producers of the cytokine interferon-γ, which is not only associated with protection against infection but is the principal diagnostic measurement in the *in vitro* interferon-γ (IFN-γ) release assay (IGRA) used to detect bTB+ cattle, CD4^+^ T cells are considered major players in anti-mycobacterial immunity (6). While other T cell subsets have received less attention, it is likely that these too play an effector and immunomodulatory role during the course of *M. bovis* infection. For example, CD8^+^ cytotoxic T cells have been shown to express IFN-γ after experimental infection with *M. bovis* but negatively impact on protection to bTB following vaccination (7, 8).

Another T cell subset, γδ T cells have attracted significant recent attention due in part to their ability to span innate and adaptive arms of the immune response but also because the full extent of their development and polyfunctionality remains enigmatic (9). γδ T cells are a diverse group of T lymphocytes that express T-cell receptor (TCR) consisting of TCR-γ and TCR-δ chains. Like classical αβ T cells, γδ T cells are an evolutionary conserved (>430 million years) lymphocyte lineage found in the immune systems of all jawed vertebrates, including humans, mice and cattle (10). An interesting feature of the γδ T cell compartment in ruminants is that unlike in humans and mice where these cells represent less than 5% of the circulating peripheral lymphocyte pool, γδ T cells constitute up to 60% of the total blood lymphocyte population in young calves and approximately 30% in adult cattle (11).

In addition to γδ TCR expression, bovine γδ T cells, express workshop cluster (WC) 1 receptor, a transmembrane glycoprotein and pathogen recognition receptor similar to human CD163c-α (12). Based on WC1 expression, γδ T cells are broadly divided into two major subsets, WC1^+^ and WC1^-^. WC1-expressing γδ T cells are further classified into two main subpopulations, WC1.1^+^ or WC1.2^+^ (13). Murine and human studies have shown how various lineages of γδ T cells can have proinflammatory (IFN-γ expression) or anti-inflammatory function (IL-10 production) and therefore it is clear that their phenotypic plasticity and development of regulatory functions could have important consequences for the ability of cattle to mount an effective anti-mycobacterial immune response against *M. bovis* infection (14). γδ T cells are of particular interest to the study of mycobacterial immunity as they have been shown to recognize *M. bovis* antigens (15, 16), secrete sentinel cytokines including IFN-γ (17, 18), and also influence the activity of other key innate cells including dendritic cells and macrophages during mycobacterial infection (19–21). One study demonstrated that circulating bovine γδ T cells spontaneously secrete the anti-inflammatory cytokine IL-10 and can inhibit proliferation of both CD4^+^ and CD8^+^ T cells, thereby documenting a major regulatory role for these cells (22).

Multiple studies now show the importance of crosstalk between innate and adaptive cells in influencing cell subtype and function, including recent studies showing an additional influence of the host microbiome on γδ T cell function (23, 24). However, the number of studies that have comprehensively assessed bovine γδ T cells under natural infection conditions are limited. Therefore, we sought to use bulk RNA-sequencing (RNA-seq) of total lymphocyte and γδ T cell subset populations from naturally infected cattle to capture the true functional status of these critical cell subsets in response to *M. bovis* infection. Our previous work showed significantly higher number of circulating γδ T cells in *M. bovis* infected cattle (25), but their functions have not yet been elucidated. In this study we hypothesized that *M. bovis* infection induces a specific γδ T cell functional phenotype that may play a role in the progression to clinical disease and that may impact on the ability of the host to clear mycobacterial infection.

## MATERIALS AND METHODS

### Experimental Animals

Male Holstein-Friesian (*Bos taurus*) cattle were used for this study (age range 18-30 months). For the infected group, animals (*n* = 12) were selected from a herd of animals naturally infected with *M. bovis* maintained at the Department of Agriculture, Food and the Marine research farm in Longtown, Co. Kildare, Ireland. The infected group animals had tested positive for bovine tuberculosis by single intradermal comparative tuberculin test (SICTT) and also by the whole blood IFN-γ release assay (IGRA, University College Dublin, Ireland). For the control group, non-infected (negative for both the SICTT and IFN-γ release assay), age and sex matched animals (*n* = 10) were selected from a herd with that were free from bTB for more than five years. All animal procedures and experimental protocols in this study were approved by the Teagasc Animal Ethics Committee (TAEC217-2019) and carried out in accordance with the relevant institutional guidelines and under license from the Irish Health Products Regulatory Authority (HPRA no. AE19132/I019).

### Blood Sampling, Processing and Cell Separation

Peripheral blood from the jugular vein of *M. bovis*-infected and control animals was collected in 10 ml Vacutainer^®^ tubes containing EDTA anticoagulant (BD Diagnostics, Oxford, UK). For cell separation, whole blood was diluted 1:1 with phosphate buffered saline (PBS) in 50-ml conical Falcon tubes (Corning, Inc., Kaiserslautern, Germany). Peripheral blood mononuclear cells (PBMCs) were isolated from buffy coat fractions using Histopaque-1077 (Sigma-Aldrich Ireland Ltd., Wicklow, Ireland) and the Leucosep system (Greiner Bio-One Ltd., Stonehouse, UK) with gradient centrifugation at 1034 × g for 25 min. PBMCs were collected and washed in PBS to remove platelets. Contaminating red blood cells were removed using cell lysis buffer (Thermo Fisher Scientific, Waltham, MA, USA).

### Preparation of Cell Suspensions, γδ T Cell Labelling and Sorting

PBMCs were washed and re-suspended in PBS containing 2% bovine serum albumin (BSA). For labelling, cells were mixed with anti-bovine γδ TCR (GB21A, IgG2b), WC1.1 (BAG25A, IgM) and WC1.2 (CACTB32A, IgG1) monoclonal antibodies (mAbs, 1 μg/2 × 10^6^ of each, Monoclonal Antibody Center, Pullman, WA, USA), incubated at 4°C for 20 min, washed and re-suspended in 2% BSA PBS. Cells were then incubated in the dark for 30-45 min on ice with goat anti-mouse fluorochrome conjugated isotype specific mAbs (Life Technologies Corporation, Invitrogen, USA), washed twice and re-suspended in 2% BSA PBS. The cells were collected, analyzed and sorted using the FACSAria Fusion Sorter (BD Biosciences, Wokingham, UK). Compensation was used to eliminate residual spectral overlaps between individual fluorochromes. Side and forward scatter area/forward scatter width parameters or characteristics were used for the identification of viable single lymphocytes and exclusion of doublets. Propidium iodide (Thermo Fisher Scientific) was used for the exclusion of dead cells. Sorted γδ T cells and subsets were identified as having viability greater than 95% and purity greater than 99%. The γδ T cells were centrifuged and cell pellets were snap frozen on dry ice first and then stored at -80°C.

### Total RNA Extraction and Library Preparation

Total RNA was extracted from the cell pellets. RNA quantity, integrity and purity were assessed using a NanoDrop™ 1000 spectrophotometer (Thermo Fisher Scientific) and an Agilent 2100 Bioanalyzer using an RNA 6000 Nano LabChip kit (Agilent Technologies Ltd., Cork, Ireland), according to the manufacturers’ instructions. All RNA samples used for transcriptomic analysis (*n* = 6 per group) had RNA integrity number (RIN) values > 8. RNA libraries were prepared from a starting quantity of 100 ng high quality RNA using the TruSeq RNA Library Prep Kit v2 (Illumina Inc., San Diego, CA, USA). All RNA samples used for transcriptomic analysis (*n* = 6 per group) had RNA integrity number (RIN) values > 8. RNA libraries were prepared from a starting quantity of 100 ng high quality RNA using the TruSeq RNA Library Prep Kit v2 (Illumina Inc., San Diego, CA, USA). Individually barcoded RNA-seq libraries were pooled in equimolar quantities and the quantity and quality of the final pooled libraries (two different pools in total) were assessed as described above. RNA-seq library sample pool construction was performed using an NEB Next® Ultra RNA Library Prep Kit for Illumina® (Cat No. 7530) and detailed in **Supplementary Table S1**. Cluster generation and high-throughput 150 bp paired-end sequencing of the pooled RNA-seq libraries were performed on a NovaSeq 6000 using an S4 flow cell prepared with the Illumina 300 cycle Reagent Kit (v1.5). All RNA-seq data generated for this study have been deposited in the ArrayExpress database under project accession number E-MTAB-13111.

### RNA-Seq Data Processing and Analysis

Sequence quality and composition were checked using FastQC (version 0.11.8) software (26). Adapter sequence reads were removed and quality trimming was carried out using the fastP adapter removal software (27). RNA seq data was processed as described previously (28). Briefly, 50 million, 2 × 150 bp paired-end sequence reads were generated from each RNA sample. Quality filtering of RNA-seq read pairs yielded a mean of 69,161,585 reads per individual library (48 libraries in total). The filtered RNA seq paired-end reads were mapped to the most recently annotated version of the *Bos taurus* reference genome (ARS-UCD1.2, GenBank assembly accession: GCA_002263795.2) (29) using the STAR aligner (version 2.7) (30). A mean of 64,133,177 read pairs (92.63%) were uniquely mapped to locations in the ARS-UCD1.2 bovine genome assembly [**Supplementary Table S1**]. Aligned reads were assigned to genomic features using featureCounts (31) and the resulting quantification files were annotated at gene level via biomaRt and GO.db (32). The annotated gene counts were then normalised and differential expression analysis performed with DESeq2 (version 1.20.0) (33). Lowly expressed reads and extreme count outliers were removed within DESeq2 using the Cook’s distance. *P* values were adjusted for multiple testing using the Benjamini-Hochberg (B-H) false discovery rate (FDR) method (34). The criteria for detection of significantly differentially expressed genes (DEGs) were an FDR-adjusted *P*-value less than 0.1 (*P*_adj._ < 0.1) and a |log_2_ FC| ≥ 1.5, which was incorporated into the statistical model in DESeq2.

### Ingenuity Pathway Analysis

The Ingenuity^®^ Pathway Analysis (IPA) software package (35) with the Ingenuity^®^ Knowledge Base (Qiagen, Redwood City, CA, USA; release date December 2022) was used to identify overrepresented (enriched) canonical pathways and interaction networks for the set of DEGs. IPA^®^ Core Analysis was performed using the default settings with the user data set as the background, high predicted confidence and all nodes selected. The significance of the association of genes with each canonical pathway was determined using a B-H adjusted right-tailed Fisher’s exact test. Gene networks (graphical representations of the molecular interactions among genes) were generated based on their relevance and connectivity with the genes in the input data. Each network was assigned a score equivalent to the negative exponent of the right-tailed Fisher’s exact test for each network. The functions of the DEGs were evaluated using the Ingenuity^®^ Knowledge Base, the GeneCards Suite (36), and the NCBI resources (37) and Uniprot (38) databases.

## RESULTS AND DISCUSSION

### Gamma-delta (γδ) T cell Subset Frequency Within Circulating Lymphocytes of BTB+ and Control Cattle

The proportions of γδ T subpopulations in the bTB+ (*n* = 12) and non-infected control (*n* = 10) cattle groups were assessed using flow cytometry on live lymphocytes in peripheral blood mononuclear cells (PBMCs). The gating strategy shows the identification of WC1.1^+^, WC1.2^+^ and WC1^-^ subpopulations within the γδ TCR^+^ subpopulation and γδ^-^ cells were identified by gating on the γδ TCR^-^ cells (**Figure 1**). No significant difference in the total γδ^+^ T cells proportions (**Figure 2A**), or WC1.1^+^ (**Figure 2B**), WC1.2^+^ (**Figure 2C**) or WC1^-^ (**Figure 2D**) subsets was detected in the *M. bovis*-infected animals compared to the non-infected controls. Significant inter-individual animal variation in all four subsets was apparent in both experimental groups (**Figure 2E**).

**Figure 1:**
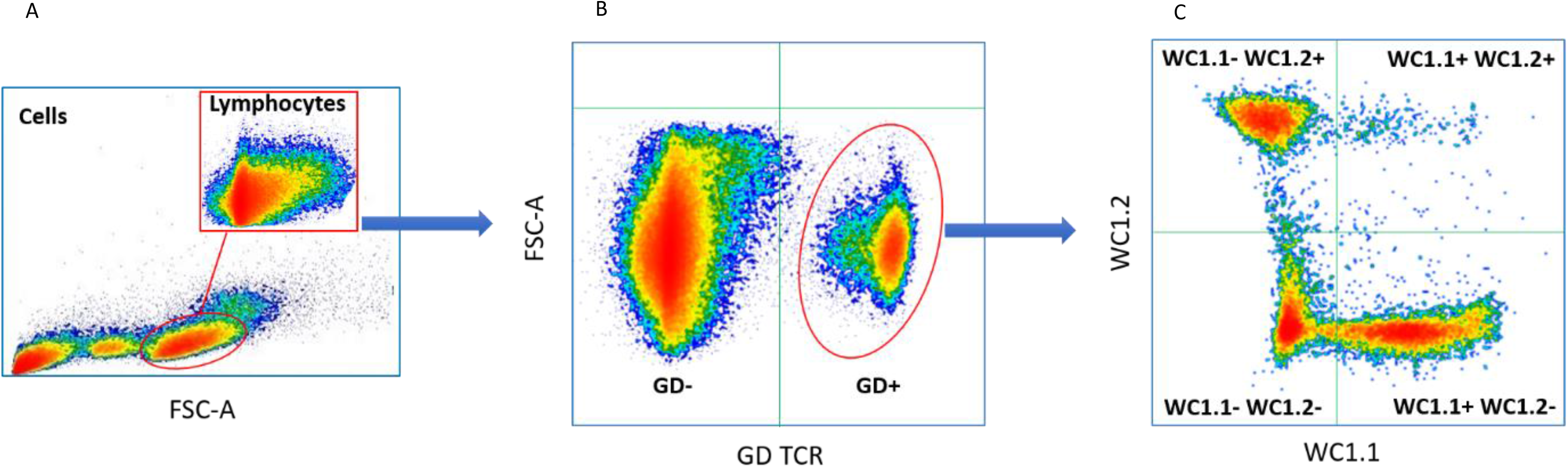
Typical gating strategy for flow cytometric γδ T cell analysis and sorting: (A) Forward and side scatter shows the gating on lymphocytes; (B) Lymphocyte separated into γδ TCR^+^ and γδ TCR^-^ sub-populations; (C) γδ TCR^+^ T cells separated into three major sub compartments based on the expression of WC1.1^+^; WC1.2^+^ or WC1^-^.

**Figure 2:**
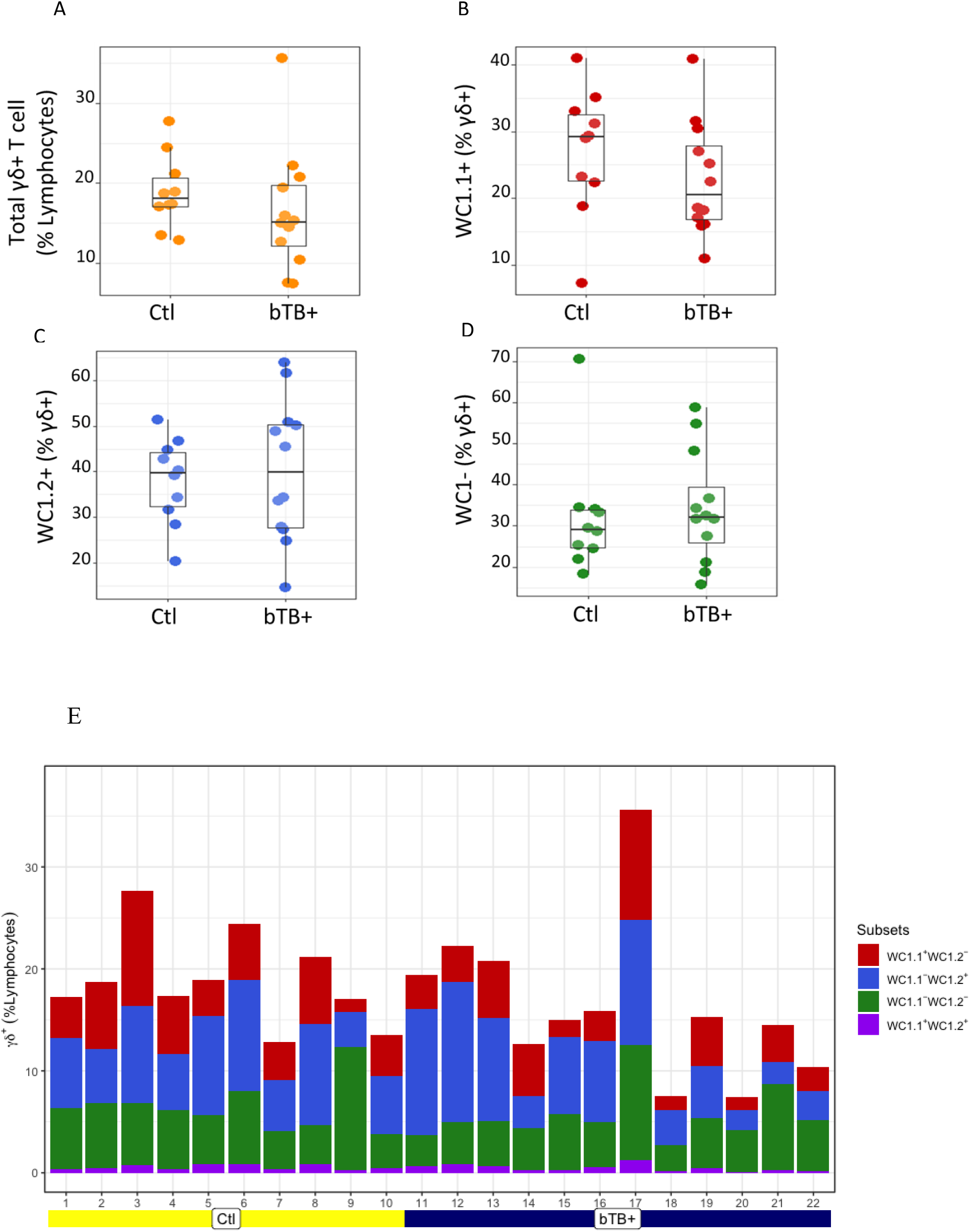
Assessment of (A) Total γδ TCR^+^; (B) WC1.1^+^; (C) WC1.2^+^ and (D) WC1^-^ subsets in bTB+ (*n* = 12) and non-infected control (*n* = 10) cattle. Data presented as standard error mean (SEM) of percentages; (E) Individual animal data for γδ T cell subsets in control and bTB+ groups, expressed as a % of lymphocytes.

### Preferential Differential Gene Expression Within WC1.1^+^ γδ T cell Compartment

Statistical analysis demonstrated that differential gene expression was evident between the bTB+ and the non-infected control cattle groups, predominantly in the WC1.1^+^ γδ T cell compartment. A total of 189 genes were observed to be differentially expressed in WC1.1^+^ γδ T cells. A smaller number of genes were differentially expressed between groups in the WC1.2^+^ γδ T cell (19 genes) and WC1^-^ γδ T cell (33 genes) compartments. Only seven genes were differentially expressed in the γδ^-^ T cells. **Supplementary Table S2** provides detailed information on the differential expression analysis results for each comparison. A Venn diagram for the significant DEGs across cell compartments between the bTB+ and non-infected control cattle groups is shown in **Figure 3**, and very minor overlap in DEGs is apparent between subsets. Most of the DE genes in the WC1.1^+^ γδ T cells were increased in expression in the bTB+ cattle (89%) and this asymmetric response is shown in the volcano plot in **Figure 4**. The range in log_2_ fold change for genes exhibiting significantly different expression in the bTB+ group ranged from – 21.83 to 29.2 (**Supplementary File S2**).

**Figure 3:**
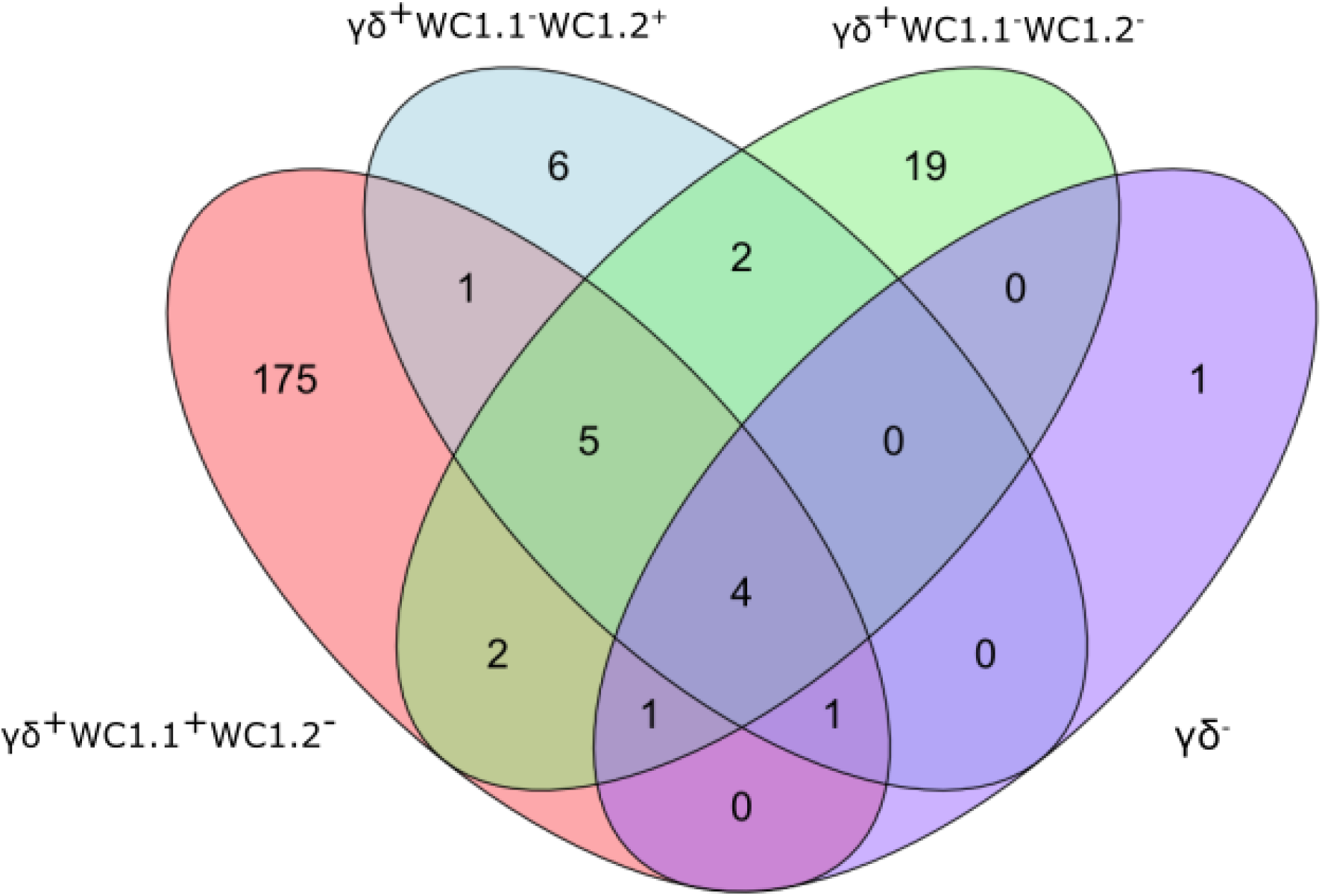
Venn Diagram showing the overlap of DEGs (FDR-*P*_adj._ < 0.10; |log_2_FC| ≥ 1.5) between γδ T cell subsets in the bTB+ group (*n* = 6) compared to the non-infected control group (*n* = 6).

**Figure 4:**
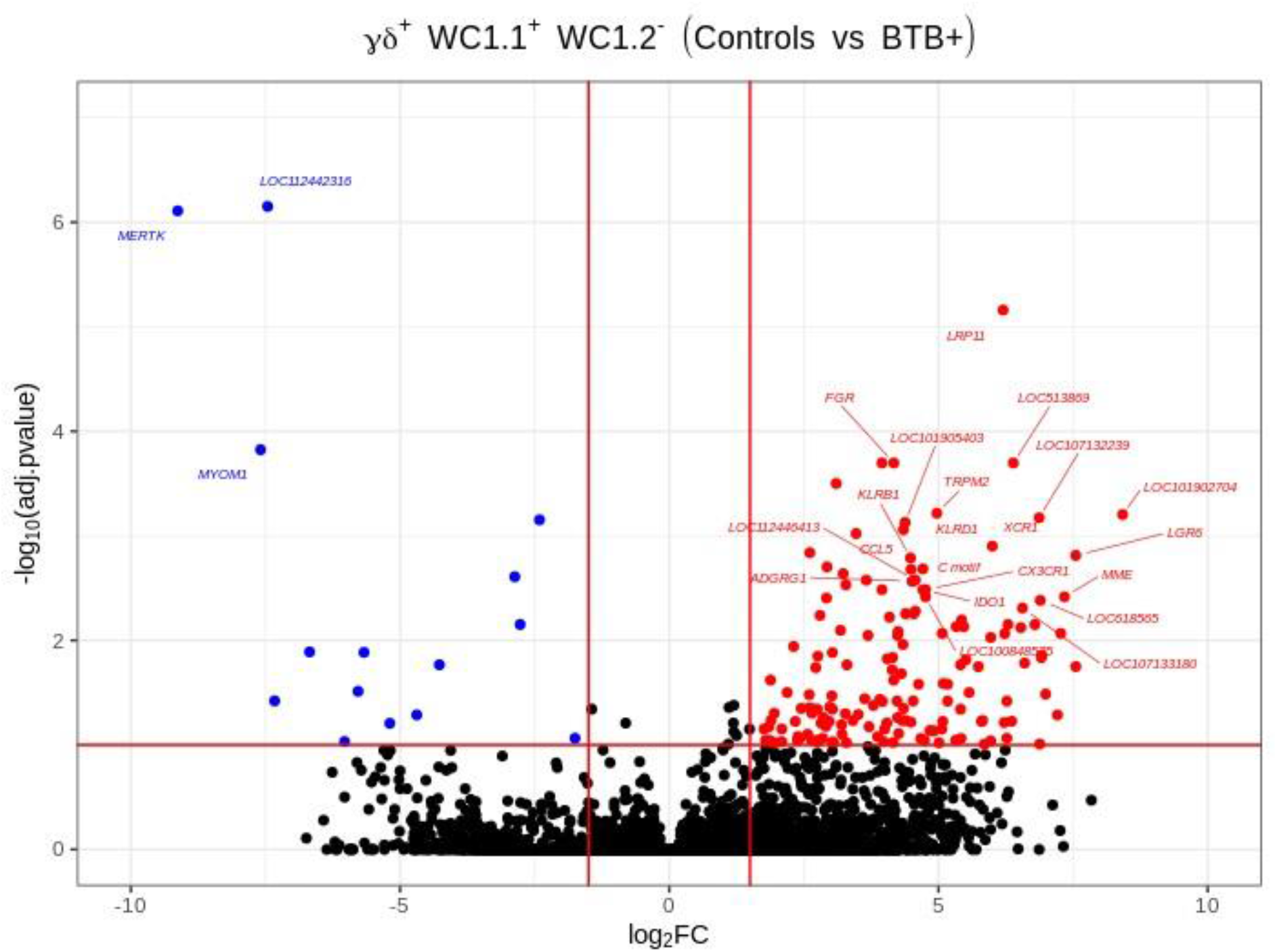
Volcano plot showing differential gene expression for the WC1.1^+^ γδ T cell subset in the bTB+ group (*n* = 6) compared to the non-infected control group (*n* = 6). The FDR-*P*_adj._ < 0.10 and |log_2_FC| ≥ 1.5 thresholds are shown as red lines.

The 168 significantly upregulated genes includes genes which encode proteins with well characterized immune roles evident from Cluster of Differentiation (CD) protein identifiers, including the cell adhesion marker CD2 (log_2_FC = 2.92; FDR-*P* = 0.0039), cell surface glycoproteins CD6 (log_2_FC = 2.75; FDR-*P* = 0.0455), CD8A (log_2_FC = 2.6; FDR-*P* = 0.0443), CD8B (log_2_FC = 2.6; FDR-*P* = 0.0330), and CD38 (log_2_FC = 3.47; FDR-*P* = 0.0010), an enzyme that functions in intracellular signalling and as a receptor with a role in endothelial adhesion (39). Interestingly, a new CD8A/B expressing subset of γδ T cells that can also express IFN-γ has been reported (40) but to-date, this subset has not been identified in domestic cattle. The *CD86* gene encoding a type I membrane protein member of the immunoglobulin superfamily that regulates T cell activation (41) was also increased in expression (log_2_FC = 2.42; FDR-*P* = 0.0867). *CD244*, a gene encoding a signalling lymphocyte activation molecule (SLAM) family immunoregulatory receptor found on many immune cell types was also upregulated (log_2_FC = 2.38; FDR-*P* = 0.0901). In addition, *CCL5*, which encodes a potent cytotoxic T cell activator (C-C motif chemokine ligand 5) (42), exhibited increased expression (log_2_FC = 4.49; FDR-*P* = 0.0021). The *CCR5* gene encoding a receptor for CCL5 was also upregulated (log_2_FC = 4.0; FDR-*P* = 0.0706), as was the *CCR7* gene (log_2_FC = 2.45; FDR-*P* = 0.0447). The CCR7 protein has been shown to regulate T cell homing and dendritic cell maturation (43). The indoleamine-2,3-dioxygenase 1 gene (*IDO1*) involved in T regulatory cell development (44) and the interleukin 2 receptor gene (*IL2RB*), which regulates T cell-mediated immune responses, were also significantly increased in expression for the bTB+ group (log_2_FC = 4.71; FDR-*P* = 0.0033; and log_2_FC = 2.31; FDR-*P* = 0.0114, respectively). A microRNA (miRNA) gene (*MIR2901*) exhibited the largest expression fold-change (log_2_FC = 23.67; FDR-*P* = 8.53 × 10^-56^); however, no functional data exists for the mir-2901 miRNA in cattle. Finally, 42 of the upregulated genes had NCBI LOC symbols for which detailed functional information is also limited (**Supplementary File 2**).

The 21 genes that exhibited decreased expression in the bTB+ group included the gamma-aminobutyric acid receptor subunit alpha-2 gene (*GABRA2*: log_2_FC = -14.14; FDR-*P* = 2.58 × 10^-6^) and the protocadherin 11 X gene (*PCDH11X*: log_2_FC = -11.09; FDR-*P* = 0.0006). *MERTK* also exhibited decreased expression (log_2_FC = -9.13; FDR-*P* = 7.78 ×10^-7^) and encodes a receptor tyrosine kinase, which is a key phagocytic receptor in the immune system that regulates many physiological processes including phagocytosis of apoptotic cells (45). Also downregulated was expression of the C-type lectin domain family 5 member A gene (*CLEC5A*), which has a role in the induction of many cytokines and chemokines thereby amplifying the innate immune response (log_2_FC = -5.67; FDR-*P* = 0.01302) (46). Interestingly, however, differential expression of cytokine genes was not a defining feature of the DEG data set. The 20 genes that exhibited the highest expression fold changes (up-and downregulated) are shown in **Table 1**.

**Table 1:** Genes exhibiting statistically significant (FDR-*P*_adj._ < 0.10; |log_2_FC| ≥ 1.5) increased and decreased expression in WC1.1+ γδ T cells for the bTB+ group (*n* = 6) compared to the non-infected control group (*n* = 6). The top ten genes ranked by fold-change are shown in each case.

| Upregulated genes |  |  |  |
| --- | --- | --- | --- |
| Gene name | Gene Symbol | Log<sub>2</sub> FC | FDR-$P_{\text{adj.}}$ |
| Epithelial membrane protein 2 | *EMP2* | 23.83 | $5.97 \times 10^{-35}$ |
| Bta-mir-2901 | *MIR2901* | 23.67 | $8.53 \times 10^{-56}$ |
| Chromosome 13 C20orf204 homolog | *C13H20orf204* | 22.61 | $1.75 \times 10^{-9}$ |
| C-type lectin domain family 7 member A-like | *LOC101902704* | 8.42 | 0.0006 |
| Leucine rich repeat containing G protein-coupled receptor 6 | *LGR6* | 7.55 | 0.0015 |
| Arachidonate 15-lipoxygenase | *ALOX15* | 7.55 | 0.0178 |
| Immunoglobulin superfamily containing leucine rich repeat | *ISLR* | 7.21 | 0.0517 |
| Mitogen-activated protein kinase kinase kinase 6 | *MAP3K6* | 6.99 | 0.0326 |
| Killer cell immunoglobulin-like receptor, three domains, long cytoplasmic tail, 1 | *KIR3DL1* | 6.92 | 0.0138 |
| Small nucleolar RNA SNORD65 | *LOC112442827* | 6.91 | 0.0146 |
| Downregulated genes |  |  |  |
| Gamma-aminobutyric acid type A receptor subunit alpha2 | *GABRA2* | -14.14 | $2.58 \times 10^{-6}$ |
| Protocadherin 11 X-linked | *PCDH11X* | -11.09 | 0.0006 |
| MER proto-oncogene, tyrosine kinase | *MERTK* | -9.13 | $7.78 \times 10^{-7}$ |
| Myomesin 1 | *MYOM1* | -7.59 | 0.0001 |
| Uncharacterized | *LOC112442316* | -7.46 | $7.06 \times 10^{-7}$ |
| Uncharacterized | *LOC101907004* | -7.33 | 0.0380 |
| Matrix metalloproteinase 19 | *MMP19* | -6.68 | 0.0129 |
| Uncharacterized | *LOC112446773* | -6.03 | 0.0930 |
| RAP1 GTPase activating protein | *RAP1GAP* | -5.78 | 0.0307 |
| C-type lectin domain containing 5A | *CLEC5A* | -5.67 | 0.0130 |

**Table 2:** The top-ranked *Diseases and Disorders* (A), *Molecular and Cellular Functions* (B) and (C) *Physiological System Development and Function* categories associated with *M. bovis* infection status identified using IPA^®^.

**A**
| **Diseases and Disorders** | **B-H *P*-value range** | **Number of genes** |
| --- | --- | --- |
| Inflammatory Response | $9.9 \times 10^{-12} - 3.0 \times 10^{-3}$ | 68 |
| Organismal Injury and Abnormalities | $9.3 \times 10^{-10} - 3.4 \times 10^{-3}$ | 107 |
| Neurological Disease | $1.5 \times 10^{-9} - 3.4 \times 10^{-3}$ | 55 |
| Inflammatory Disease | $1.0 \times 10^{-8} - 2.6 \times 10^{-3}$ | 59 |
| Connective Tissue Disorders | $5.9 \times 10^{-8} - 7.8 \times 10^{-5}$ | 42 |

**B**
| **MOLECULAR AND CELLULAR FUNCTIONS** | **B-H *P*-VALUE range** | **NUMBER OF GENES** |
| --- | --- | --- |
| Cell-to-Cell Signalling and Interaction | $8.5 \times 10^{-12} - 3.3 \times 10^{-3}$ | 52 |
| Cell Death and Survival | $9.9 \times 10^{-12} - 2.3 \times 10^{-3}$ | 55 |
| Cellular Compromise | $1.0 \times 10^{-11} - 2.3 \times 10^{-3}$ | 26 |
| Cellular Development | $1.2 \times 10^{-10} - 3.4 \times 10^{-3}$ | 35 |
| Cellular Growth and Proliferation | $1.2 \times 10^{-10} - 3.4 \times 10^{-3}$ | 39 |

**C**
| **Physiological System Development and Function** | **B-H *P*-value range** | **Number of genes** |
| --- | --- | --- |
| Haematological System Development and Function | $8.5 \times 10^{-12} - 3.4 \times 10^{-3}$ | 55 |
| Immune Cell Trafficking | $9.9 \times 10^{-12} - 3.4 \times 10^{-3}$ | 47 |
| Lymphoid Tissue Structure and Development | $1.2 \times 10^{-10} - 3.4 \times 10^{-3}$ | 40 |
| Tissue Morphology | $1.6 \times 10^{-8} - 3.0 \times 10^{-3}$ | 44 |
| Cell-mediated Immune Response | $3.3 \times 10^{-8} - 3.4 \times 10^{-3}$ | 31 |

### Gene Network and Pathway Analysis of WC1.1+ γδ T Cells

The WC1.1^+^ γδ T cell DEG data set was further analysed using IPA^®^ to identify enriched canonical pathways and biological interaction networks. Out of the 189 significant DEGs, 125 analysis-ready genes could be mapped to genes in the Ingenuity^®^ Knowledge Base, of which 111 had increased expression and 14 had decreased expression, against a background set of 14,109 analysis-ready genes (7857 with increased expression, 5876 with decreased expression and 376 with no change in expression) from the 18,310 detectable genes that were mapped by IPA^®^. Canonical pathway analysis revealed that the DEGs were enriched in several notable signalling pathways (**Figure 5A**). The top biological pathways were *Natural Killer Cell Signaling*, *Crosstalk between Dendritic Cells and Natural Killer Cells*, *Th1 and Th2 Activation Pathway*, *CTLA4 Signaling in Cytotoxic T Lymphocytes*, and the *CREB Signaling in Neurons* represented by the DEGs shown in **Figure 5B**.

**Figure 5:**
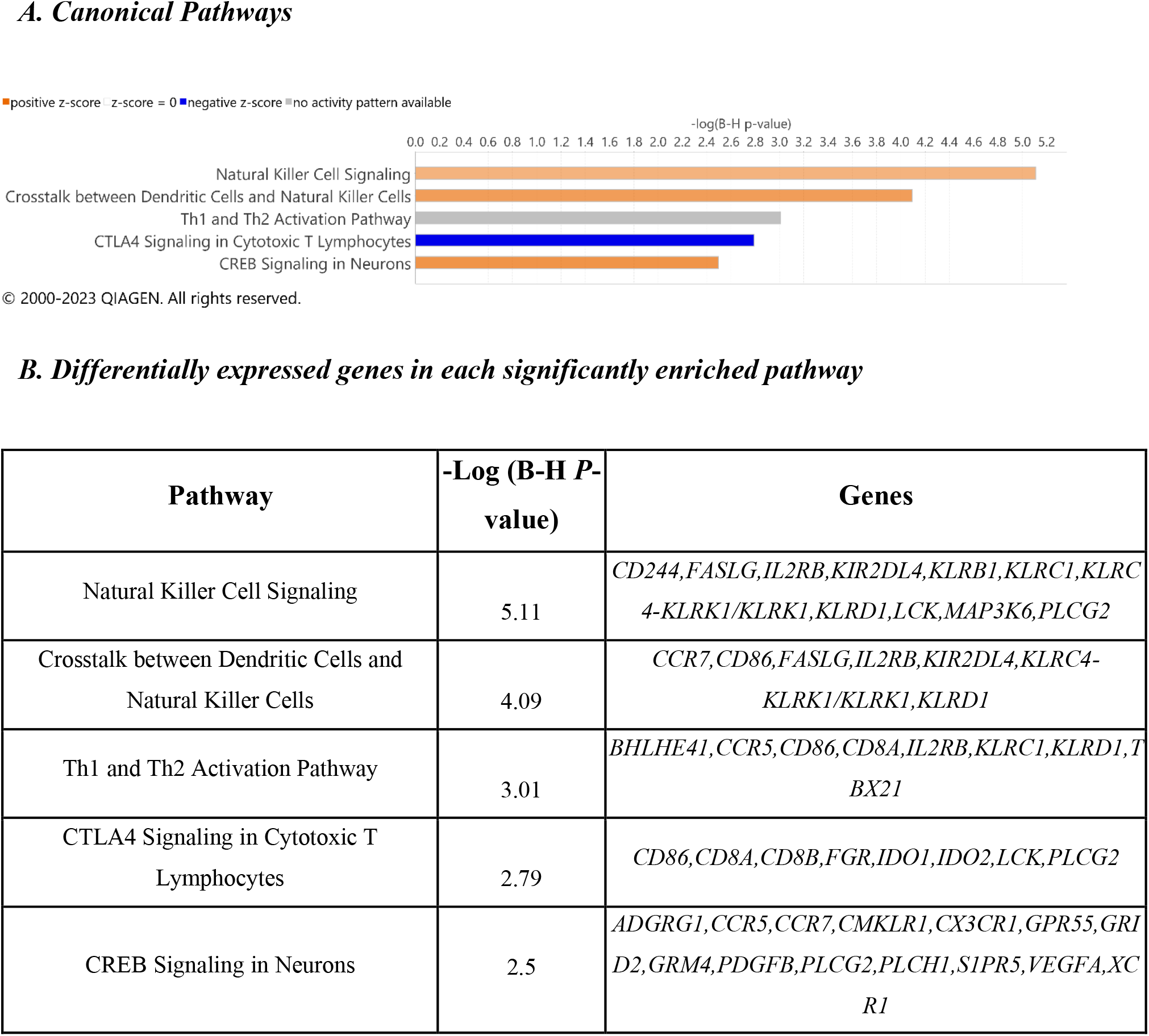
Canonical pathway enrichment results from IPA^®^ showing (A) Top five biological pathways associated with *M. bovis* infection in cattle IPA^®^; (B) The DEGs present within each enriched pathway.

The top diseases and biological functions included *Inflammatory Response, Organismal Injury and Abnormalities, Cell-to-Cell Signalling and Interaction, Cell Death and Survival, Hematological System Development and Function* and *Immune Cell Trafficking* (see **Tables 2A**, **2B** and **2C**). Eleven biological interaction networks were generated from the 125 DEGs set using the IPA^®^ Knowledge Base (**Supplementary Table S3**). The network analysis showed that the *M. bovis* infection in cattle affected diverse biological functions and cellular processes (**Supplementary Table S4 and S5**). **Figure 6** shows the highest-ranked biological network (with a network score of 33 and 17 focus molecules), which includes *ACER2, ADGRG1, Calmodulin, CCR5, CCR7, CEMIP2, CMKLR1, CX3CR1, Erm, Fc gamma receptor, Focal adhesion kinase, G protein alpha i, Gpcr, GPR55, GRM4, Integrin, NFkB (complex), Nr1h, OSCAR, p85 (pik3r), Pka, PLC, PLC gamma, PLCG2, PLCH1, PPEF1, PPIF, PTCH1, Rac, RAS, S1PR5, Sfk, SRC (family), STAT* and *XCR1*.

**Figure 6:**
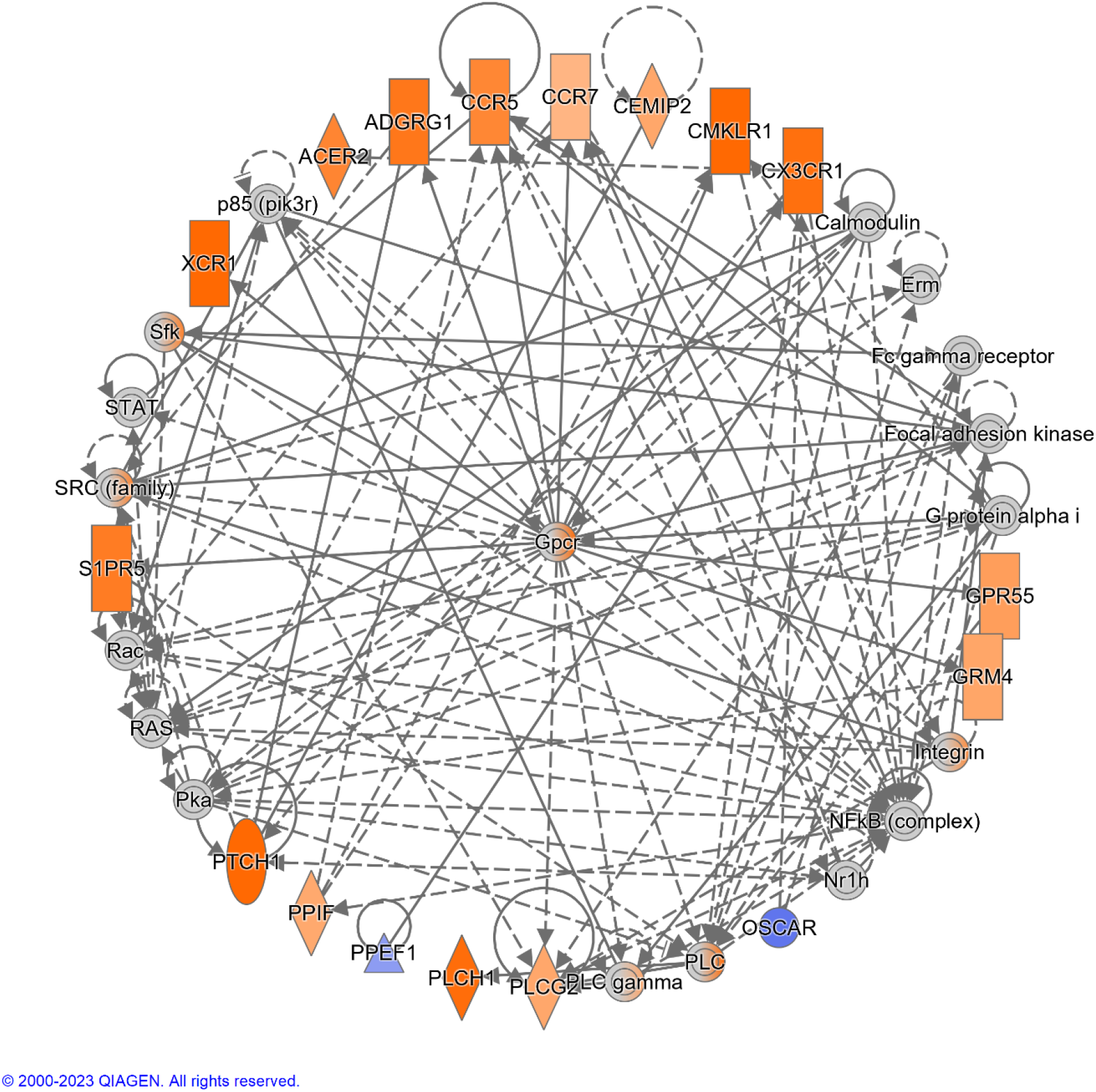
The top-ranked network generated using IPA^®^, which consisted of 17 focus genes (Network Score = 33). The top Diseases and Functions for this network are *Cell-mediated Immune Response* (*P*-value = 3.8 × 10^-7^)*, Cell-To-Cell Signaling and Interaction* (*P*-value = 7.3 × 10^-7^) and *Cellular Movement* (*P*-value = 1.0 × 10^-6^). Increased and decreased gene expression is indicated with the orange and blue colour scales, respectively. Genes that were not detectable are indicated in grey. A full legend is available at: https://qiagen.my.salesforce-sites.com/KnowledgeBase/articles/Basic_Technical_Q_A/Legend.

Despite no apparent difference in overall γδ T cell or subset numbers between the bTB+ and control groups, our results clearly show preferential activation of the WC1.1^+^ γδ T cell compartment in bTB+ infected cattle. It is surprising that of all the direct comparisons between cell types from the bTB+ and healthy, control cattle, numbers of DEG are low. This includes the examination of the γδ TCR^-^ lymphocytes, suggesting that a differential response is only maintained in the WC1.1^+^ γδ T cell compartment, even in the absence of exogenous antigen stimulation. Other studies have found that this γδ T cell subset is more inflammatory (47) and it is the WC1.1^+^ subset that localize to the site of infection after BCG vaccination (48), thereby supporting the immunoreactive ability to mycobacterial antigens of these cells.

### Differential Expression of NK-related Genes with Cytotoxic Function

One of the defining features of our analysis was the overrepresentation of genes that encode markers more often associated with the function of natural killer (NK) cells. NK cells are cytotoxic lymphocytes endowed with multiple mechanisms for killing infected cells, including those infected with mycobacteria (49). Cytotoxic cells can directly kill intracellular bacteria through granulysin-mediated delivery of granzymes (50) and these molecules have documented antimycobacterial efficacy against *M. tuberculosis* (51).

In the DEG data set generated for the present study, *CD244*, encoding a cell surface receptor expressed on natural killer (NK) cells and other T cells is upregulated in bTB+ cattle (log_2_FC = 2.38; FDR-*P* = 0.0901). The CD244 protein modulates cellular cytotoxicity and plays an important role in regulating production of IFN-γ as well as both CD4^+^ and CD8^+^ T cell immunity during TB disease (52, 53). The natural killer cell granule protein 7 gene (*NKG7*), a regulator of lymphocyte granule exocytosis and inflammation, also exhibited increased expression (log_2_FC = 2.07; FDR-*P* = 0.0929). NK receptors also play a direct role in the regulation of γδ T-cell-mediated immune responses, likely reflecting important cellular crosstalk in this context, and their activation leads to the release of cytotoxic granules containing granzymes (54). Also of relevance is the differential expression of granzyme A and M genes (*GZMA* and *GZMM*), which have documented antimycobacterial activity (log_2_FC = 4.04; FDR-*P* = 0.0618 and log_2_FC = 4.25; FDR-*P* = 0.0082, respectively), have been previously reported to be expressed in the γδ T cells of other species (55, 56). In conjunction with the granzyme H gene (currently annotated to *LOC788601;* log_2_FC = 6.88; FDR-*P* = 0.0980), these genes all exhibit increased expression in bTB+ cattle, suggesting a role in the immune response to *M. bovis* infection.

The eomesodermin gene (*EOMES*) encodes a transcription factor with a crucial role in regulating cytotoxic function, development, and survival of a range of immune cells such as NK and CD8^+^ T cells (57–59). In addition, the T-box transcription factor 21 gene (*TBX21*) encodes a transcription factor that plays a central role in CD4^+^ Th1 lineage development (60, 61). Consequently, *EOMES* and *TBX21* encode proteins that control the development and function of a repertoire of cells in the innate and adaptive compartments of the immune system (59). Both *EOMES* and *TBX21* showed increased expression in bTB+ cattle (log_2_FC = 4.35; FDR-*P* = 0.0447 and log_2_FC = 3.95; FDR-*P* = 0.0002, respectively). The genes currently annotated to *LOC104968634* and *LOC112448791* were also upregulated (log_2_FC = 4.14; FDR-*P* = 0.0191 and log_2_FC = 4.57; FDR-*P* = 0.0052, respectively) and encode the NK2B and granulysin (GNLY) antimicrobial peptides, respectively. *LOC104968634* has also been shown to be upregulated in bovine alveolar macrophages challenged *in vitro* with *M. bovis* (62). The differential gene expression for the DEGs with cytotoxic NK-related functions is shown in **Figure 7**.

**Figure 7:**
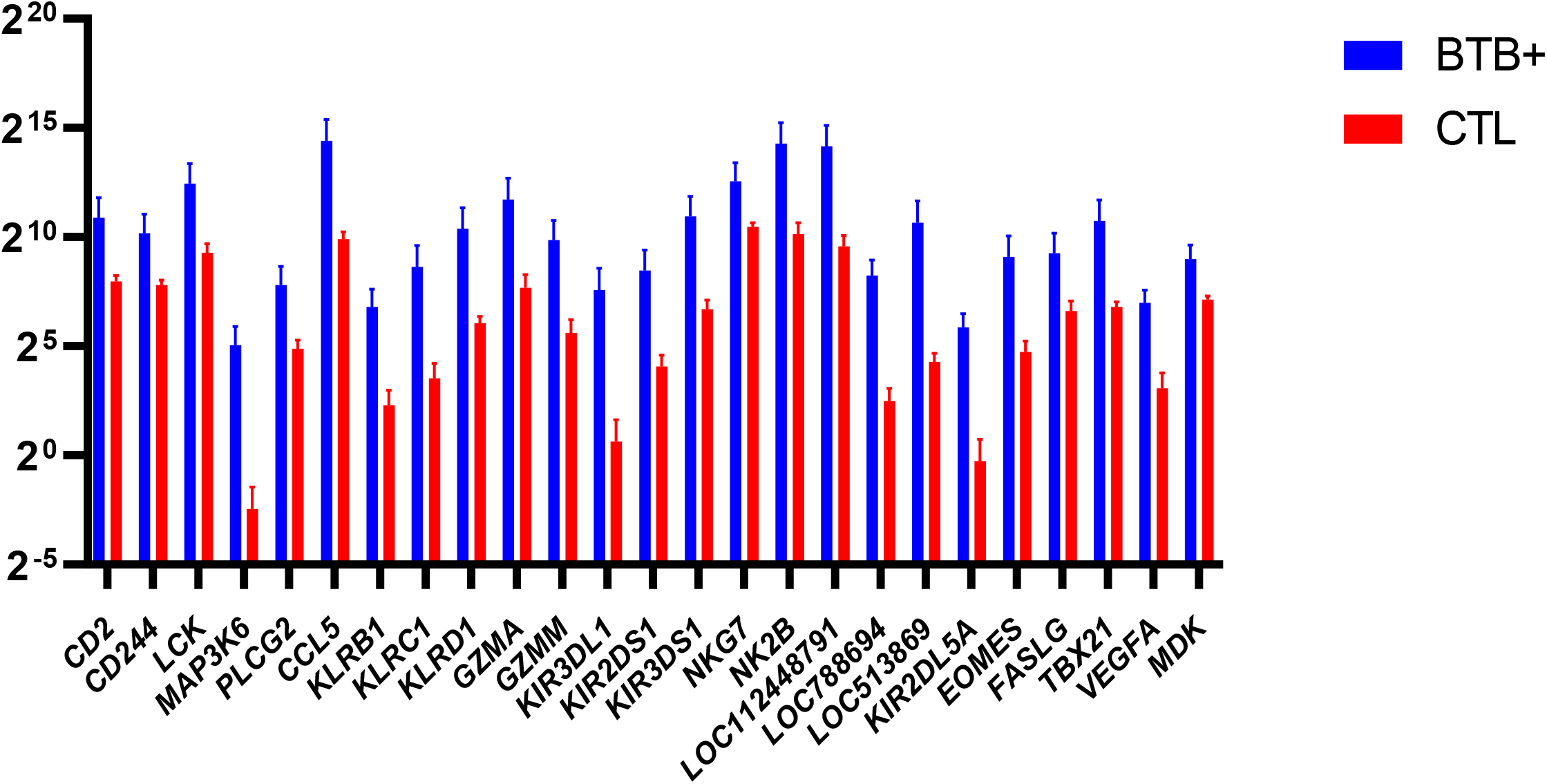
Natural Killer (NK) related genes identified as differentially expressed (FDR-*P*_adj._ < 0.10; |log_2_FC| ≥ 1.5) in WC1.1^+^ γδ T cells from the bTB+ group (*n* = 6) compared to the non-infected control group (*n* = 6). The log_2_ normalized read counts ± SEM are shown.

Other NK-related genes that exhibited increased expression in bTB+ cattle included four members of the killer cell lectin like receptor (KLR) family of immune inhibitory receptor genes (*KLRB1*, *KLRC1*, *KLRD1*, and *KLRK1*). *KLRB1* (log_2_FC = 4.48; FDR-*P* = 0.0016) encodes a protein that has been reported to have both costimulatory and coinhibitory roles in T-cells and is attracting significant attention with other KLR proteins for their potent therapeutic potential (63, 64). *KLRD1* (log_2_FC = 4.35; FDR-*P* = 0.0009) encodes a protein that has been shown to induce functional exhaustion of cytotoxic NK and CD8^+^ T cells (65–67). *KLRK1* (log_2_FC = 3.3; FDR-*P* = 0.0171) encodes the natural killer group 2D protein (NKG2D), which has a documented roles in immunoprotection from infection via γδ T cell activation (68), and pulmonary clearance of bacteria (69). Multiple killer cell immunoglobulin like receptor (KIR) genes were also upregulated in the present study, including *KIR2DL5A, KIR2DS1, KIR3DL1*, and *KIR3DS1.* Individual KIR genes as well as specific haplotypes has been previously associated with resilience to TB disease in human populations (70, 71).

The association of some of these NK-related functions has only been superficially investigated in γδ T cells previously and has not been documented in cattle naturally infected with *M. bovis* indicating potential relevance to bTB disease. These findings are supported by related research in murine models. Whereas γδ TCR knockout mice support a protective role for γδ T cells in mycobacterial infection (72), control of mycobacterial infection was seriously impaired in NK cell knockout mice (73), suggesting that the adoption of these functions by γδ T cells is a critical disease control measure. It is of further relevance that depletion of the NK cell subset has been reported in human patients with TB (74) and the authors report that this subset is GZMB^+^, therefore indicating a potential loss of granzyme function, at least from the NK cell subset associated with the progression to clinical disease. It is therefore plausible that γδ T cell granzyme expression may partially compensate for this loss of anti-mycobacterial function. However, comprehensive characterization of the NK cell subset has not been performed in *M. bovis*-infected cattle to date but based on the results described here, this avenue warrants further investigation.

### Minor Gene Expression Differences in WC1.2^+^, WC1^-^ γδ^+^ and γδ^-^ T Cell Compartments

Smaller numbers of significant DEGs were detected between groups in the other cellular compartments. In the WC1.2^+^ γδ T cell, 15 DEGs were identified between the bTB+ and non-infected control cattle groups. Six genes were upregulated, and nine genes were downregulated in the bTB+ group, respectively. Genes encoding cytochrome 8B (*COX8B*) (log_2_FC = 18.9; FDR-*P* = 2.41 × 10^-16^), cytochrome P450 family 17 subfamily A member 1 (*CYP17A1*; annotated to *LOC112444495*) (log_2_FC = 28.38; FDR-*P* = 0.0025), and LDL receptor related protein 11 (*LRP11*) (log_2_FC = 5.31; FDR-*P* = 0.0012) were increased in expression. The myomesin 1 gene (*MYOM1*), the protocadherin 11 X-linked gene (*PCDH11X*), and a solute carrier family 22 member 9-like gene (annotated to *LOC517475*) were decreased in expression.

Of the 28 DEGs in the WC1^-^ γδ T cell compartment, the majority (64%) are also decreased in expression in the bTB+ cattle. Some of the transcripts are differentially expressed in the same manner as the WC1.2^+^ subset. These include upregulation of the cytochrome P450 17A1 gene (*LOC112444495*; log_2_FC = 28.25; FDR-*P* = 0.0019) and the LDL receptor related protein 11 gene (*LRP11*; log_2_FC = 4.47; FDR-*P* = 0.0198), and downregulation of the myomesin 1 gene (*MYOM1*; log_2_FC = -12.65; FDR-*P* = 1.44 × 10^-10^) and the protocadherin 11 X-linked gene (*PCDH11X*; log_2_FC = -10.32; FDR-*P* = 0.0040) indicating differential expression of these genes is not specific to particular cell subsets. Downregulation of additional genes including the aldehyde oxidase gene (*AOX1*; log_2_FC = -20.04; FDR-*P* = 1.10 × 10^-8^) was also detected. Aldehyde oxidase catalyses the production of hydrogen peroxide and can also catalyse the formation of superoxide (75). which may impact production of ROS species in these cells.

Only six DEGs were detected in the γδ-T cells, all of which were decreased in expression in the bTB+ cattle group. Consistent decreased expression in *PCDH11X* (log_2_FC = -11.46; FDR-*P* = 0.0037) and *MYOM1* (log_2_FC = -8.81; FDR-*P* value 3.56 × 10^-5^) was apparent in these lymphocytes. A full list of DEGs detected across the different biological contrasts is provided in **Supplementary Table S2.**

## CONCLUSIONS

The eradication of bTB is a major policy aim in countries where the disease is endemic as it has widespread negative impacts on the agri-food sector and poses a zoonotic risk to human health (3). A comprehensive understanding of all aspects of the host-pathogen relationship are required to better understand immune mechanisms associated with pathogenesis and to improve diagnostic sensitivity and vaccination performance (76).

Gamma-delta (γδ) T cells connect the innate and adaptive arms of the immune response with known anti-mycobacterial function but although expanded in cattle, knowledge of their functional capacity in livestock species remains limited. Here we document significant differential expression indicating specific activation of WC1.1^+^ γδ T cells in response to *M. bovis* infection. We have identified differential regulation of a number of NK-related cytotoxic functions, which may be important for disease pathogenesis and that shed new light on the functional capacity of bovine γδ T cells under natural infection conditions.

## DATA AVAILABILITY STATEMENT

All RNA-seq data generated for this study have been deposited in the ArrayExpress database under project accession number E-MTAB-13111.

## ETHICS STATEMENT

All experimental procedures involving animals were conducted under ethical approval and experimental license (HPRA No. AE19132/I019) from the Irish Health Products Regulatory Authority (HPRA) in accordance with the Cruelty to Animals Act 1876 and in agreement with the European Union (Protection of Animals Used for Scientific Purposes) regulations 2012 (S.I. No. 543 of 2012).

## AUTHOR CONTRIBUTIONS

Conceived the study: KGM. Performed experiments and interpreted data: SAB, ME, KGM. Performed the bioinformatics analysis: TH and CR. Wrote the manuscript: KGM, SAB and DEM. All authors reviewed and approved the final manuscript.

## ACKNOWLEDGEMENTS

The authors would like to express their gratitude to Edward Mulligan, Colm Brady and the staff at the Department of Agriculture, Food, and the Marine (DAFM) farm in Longtown, Kildare, Ireland for their help with sample collection. In addition, we would like to thank Barry Moran (Trinity College Dublin) for his expertise with flow cytometry and Dr Alia Parveen (University College Dublin) for her assistance with data file submission.

## SUPPLEMENTARY MATERIAL

The Supplementary Material for this article can be found online at XXXX.

## DECLARATIONS

### CONSENT FOR PUBLICATION

Not applicable.

