## Supplementary material for "Preferential differential gene expression within the WC1.1^+^ γδ T cell compartment in cattle naturally infected with *Mycobacterium bovis*"

**Note:** British English language style preferred for publication.

**Supplementary Tables**

**Table S1**: Filtering and mapping statistics for 48 RNA-seq libraries from *M. bovis*-infected and non-infected control animals (Contained in Excel file – **Supplementary Information** File 1). **Table S2**: Differentially expressed genes (DEGs) in the γδ T cell subsets from *M. bovis*-infected and non-infected control animals (statistically significant DEGs (included are those which had RNA seq reads ≥ 3 animals and showed ≥ ± 1.5-fold change in expression with FDR *p* adj < 0.1) shown. (Contained in Excel file – **Supplementary Information** File 2).

**Table S3**: Canonical pathways identified using IPA^®^ for RNA-seq DE gene results in the WC1.1+ γδ T cells from *M. bovis*-infected and non-infected control animals (statistically significant pathways (adjusted *P* ≤ 0.1) shown and ranked according to B-H adjusted *P*-value [smallest to largest]). (Contained in Excel file – **Supplementary Information** File 3).

**Table S4**: Biological networks identified using IPA^®^ for RNA-seq DE gene results in the WC1.1+ γδ T cells from *M. bovis*-infected and non-infected control animals (statistically significant networks (adjusted *P* ≤ 0.1) shown and ranked or scored according to the connectivity and interactions between the genes in the dataset [largest to smallest]). (Contained in Excel file – **Supplementary Information** File 4).

**Table S5**: Diseases and biological (cellular and molecular) functions identified using IPA^®^ for RNA-seq DE gene results in the WC1.1+ γδ T cells from *M. bovis*-infected and non-infected control animals (statistically significant (adjusted *P* ≤ 0.1) shown and ranked according to B-H adjusted *P*-value [smallest to largest]). (Contained in Excel file – **Supplementary Information** File 5).
